# Proactive interhemispheric disinhibition supports response preparation during selective stopping

**DOI:** 10.1101/2022.09.08.507205

**Authors:** Corey G. Wadsley, John Cirillo, Arne Nieuwenhuys, Winston D. Byblow

## Abstract

Response inhibition is essential for terminating inappropriate actions. A substantial delay may occur in the response of the non-stopped effector when only part of a multi-effector action is terminated. This *stopping-interference effect* has been attributed to nonselective response inhibition processes and can be reduced with proactive cueing. This study aimed to elucidate the role of interhemispheric primary motor cortex (M1-M1) influences during selective stopping with proactive cueing. We hypothesized that stopping-interference would be reduced as stopping certainty increased, owing to proactive recruitment of interhemispheric facilitation or interhemispheric inhibition when cued to respond or stop, respectively. Twenty-three healthy human participants performed a bimanual anticipatory response inhibition paradigm with cues signaling the likelihood of a stop-signal occurring. Dual-coil transcranial magnetic stimulation was used to determine corticomotor excitability (CME), interhemispheric inhibition (IHI), and interhemispheric facilitation (IHF) in the left hand at rest and during response preparation. Response times slowed and stopping-interference decreased with cues signaling increased stopping certainty. Proactive response inhibition was marked by a reduced rate of rise and faster cancel time in electromyographical bursts during stopping. There was a nonselective release of IHI but not CME from rest to in-task response preparation, while IHF was not observed in either context. An effector-specific CME but not IHF or IHI reduction was observed when the left hand was cued to stop. These findings indicate that the stopping-interference effect can be reduced through proactive suppression. Interhemispheric M1-M1 channels modulate inhibitory tone that supports responding, but not selective stopping, in a proactive response inhibition context.

**Significance statement:** Response inhibition is essential for terminating inappropriate actions and, in some cases, may be required for only part of a multi-effector action. The present study examined interhemispheric influences between the primary motor cortices during selective stopping with proactive cueing. Stopping selectivity was greater with increased stopping certainty and marked by proactive response inhibition of the hand cued to stop. Inhibitory interhemispheric influences were released during response preparation but were not affected by proactive cueing. These findings indicate that between-hand stopping can be selective with proactive cueing, but cue-related improvements are unlikely to reflect advance engagement of interhemispheric influences between primary motor cortices.

## Introduction

Actions often require the coordination of multiple effectors. For example, steering, accelerating, and braking require precise coordination while driving. Response-selective stopping (hereby referred to as “selective stopping”) refers to scenarios where only a subcomponent of a multicomponent action must be terminated (Wadsley et al., 2022b). Terminating an inappropriate action, *response inhibition*, supports selective stopping. However, a substantial delay occurs in the response of non-stopped effectors during selective stopping (Aron & Verbruggen, 2008; Coxon et al., 2007). This *stopping-interference effect* is a demonstrable constraint of selective stopping, which may arise from fast-acting global response inhibition (Raud et al., 2020; Wadsley et al., 2022a).

Selective stopping can be required with or without foreknowledge. For example, suddenly needing to cancel a lane change is more likely on a busy motorway than on a quiet street. Proactive response inhibition reflects inhibitory processes that occur in anticipation of a need to stop (Verbruggen & Logan, 2009). The influence of foreknowledge during selective stopping can be assessed with uninformative (reactive) and informative (proactive) warning cues (Cirillo et al., 2018; Raud & Huster, 2017). Informative cues allow for stopping to be prepared for a specified subcomponent while others can respond with certainty (e.g., “maybe stop-left” cue), whereas the forthcoming stopping requirement is ambiguous during uninformative cues (e.g., “maybe stop either” cue).

The stopping-interference effect is reduced but not abolished with informative cues. Greater selectivity is driven, in-part, by proactive response inhibition during action preparation (Majid et al., 2012). Behaviorally, response times are slowed in effectors cued to stop. Transcranial magnetic stimulation (TMS) of primary motor cortex (M1) showed a concomitant suppression of corticomotor excitability (CME) during behavioral slowing (Cai et al., 2011; Majid et al., 2012). Paired-pulse TMS studies have shown a decrease in gamma-aminobutyric acid (GABA) receptor-mediated intracortical inhibition toward effectors more likely to respond (Cirillo et al., 2018; Cowie et al., 2016). Thus, improvements in selective stopping are driven through proactive control over M1 output during action preparation.

Interhemispheric M1-M1 influences support action preparation and can be investigated with dual-coil TMS. Interhemispheric inhibition (IHI) is produced through transcallosal activation of inhibitory interneurons and is mediated by GABA-B postsynaptic receptors at long interstimulus intervals (Daskalakis et al., 2002; Irlbacher et al., 2007). Interhemispheric facilitation (IHF) is mediated by cortico-cortical interactions when probed at short (4 – 8 ms) interstimulus intervals (Baumer et al., 2006; Koch et al., 2006). Modulation of IHF and IHI occurs during action preparation and is influenced by task dynamics (e.g., Neige et al., 2021). IHI is increased in the stopping effector and released in the responding effector during reactive selective stopping (MacDonald et al., 2021). However, it is unclear how facilitatory and inhibitory M1-M1 mechanisms influence proactive response inhibition. An up-regulation of GABA-B receptor-mediated inhibition examined within M1 indicates a general role in setting an inhibitory tone that can influence the selectivity of stopping (Cirillo et al., 2018; Cowie et al., 2016). While IHF may contribute to action reprogramming, (Mars et al., 2009) it remains to be determined if IHF plays a role in selective stopping.

The present study investigated M1-M1 interhemispheric influences for selective stopping with and without proactive cueing. Selective stopping was assessed in a bimanual anticipatory response inhibition (ARI) paradigm. Dual-coil TMS positioned over bilateral M1s was used to assess IHF and IHI from motor evoked potentials during response preparation. Additional measures of selective stopping were obtained from electromyography recorded from both the stopping and responding hand. Three hypotheses were tested: 1) Stopping will be faster and stopping-interference less as stopping certainty increases. 2) IHI will be evident at rest and released during action preparation in a task-relevant effector. 3) An effector-specific upregulation of IHF will occur during informative but not uninformative cues.

## Methods

### Participants

Twenty-four healthy adults volunteered to participate. One participant was excluded due to left-handedness. The remaining 23 participants (13 female and 10 male; mean age 27.2 yrs., range 22 to 39 yr.) were all right-handed (mean laterality quotient 0.83, range 0.25 to 1; Veale, 2014). The target sample size was selected based on similar dual-coil TMS studies that have investigated selective stopping (MacDonald et al., 2021). The study was approved by the University of Auckland Human Participants Ethics Committee (Ref. UAHPEC22709).

### Task protocol

A multicomponent ARI paradigm was used to assess selective stopping (Wadsley et al., 2022b). The task was programmed in PsychoPy (v2020.2.4; Peirce et al., 2019) and interfaced with a custom Arduino Leonardo response board. Participants were seated comfortably in front of an LG 24GL600F-B monitor (144 Hz refresh rate, ∼60 cm viewing distance). The left and right index fingers rested on blocks with mechanical switches positioned ∼1 cm above such that responses required sagittal index finger abduction (Figure 1A). Switch height was adjusted to minimise postural muscle activity observed from electromyography (EMG) of first dorsal interosseous (FDI) muscles bilaterally. The display consisted of two white bars (15 cm high, 1.5 cm wide) on a grey background. A black horizontal target line was positioned behind each bar at 80% of its total height. Trial onset occurred when the bars appeared to “fill” (i.e., gradually turning black from bottom to top). Pressing the left or right switch during a trial (1.5 s) would cause the corresponding bar to cease filling.

**Figure 1:**
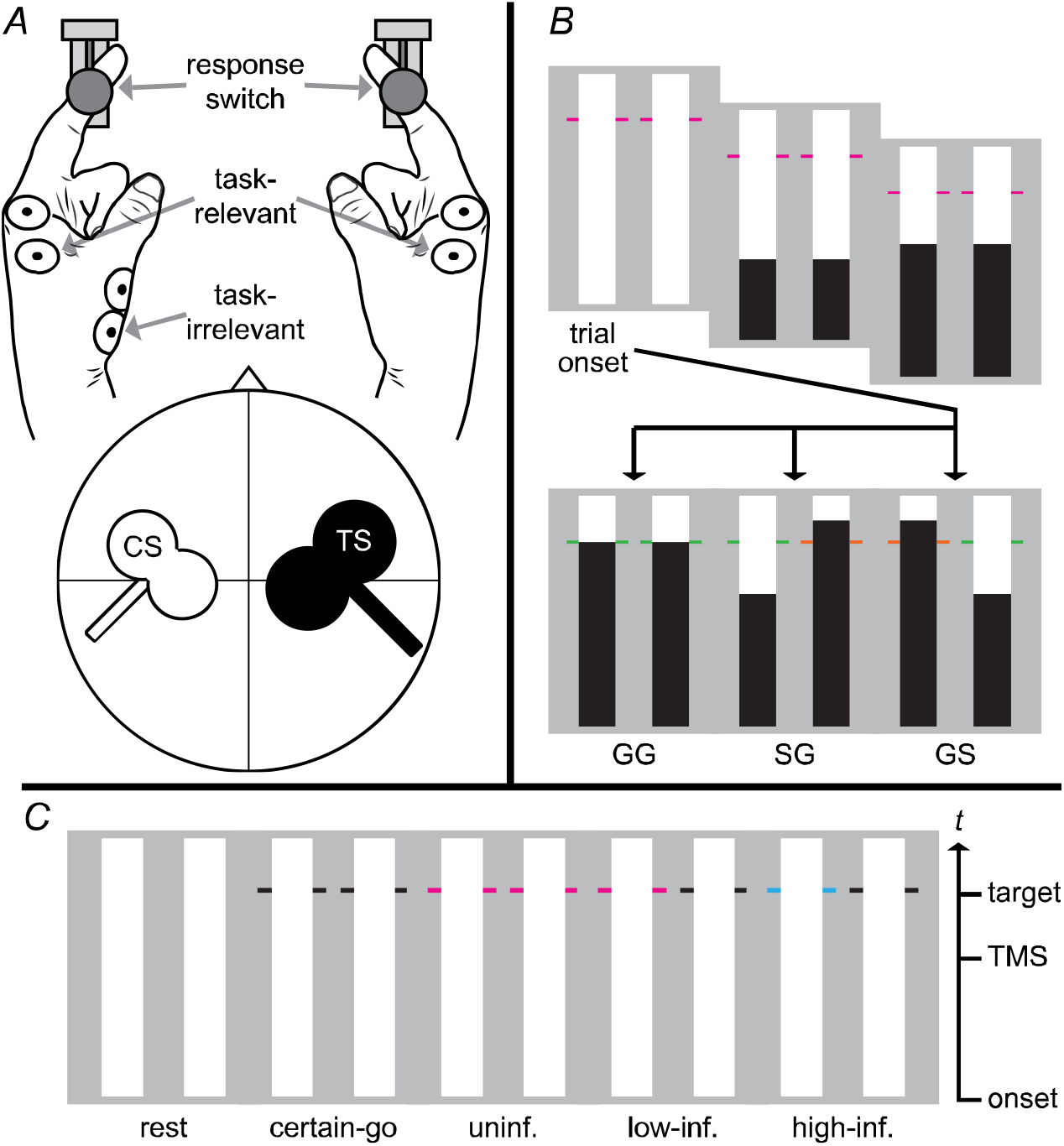
Bimanual anticipatory response inhibition paradigm. *A*: Responses were made using the left and right index fingers. The test stimulus (TS) and conditioning stimulus (CS) coils targeted the left and right hand, respectively. *B*: Timeline from trial onset. All trials began with two empty bars. The objective during go trials (GG) was to press the switches when the indicators reached the target (800 ms). During partial-stop trials either the left (stop-left go-right; SG) or right (go-left stop-right; GS) bar automatically stopped filling before the target, thus requiring the response in the corresponding hand to be cancelled. *C*: Cues were presented before trial onset to assess responses in rest, certain-go, uninformative, low-informative, and high-informative contexts. In-task stimulation was delivered 550 ms from trial onset (TMS). Stop cues were equally distributed across left and right (stop-left informative cues depicted in the figure).

The objective of most trials was to cease the bars from filling as close as possible to the target lines (go-left go-right: GG). Each bar took 1 second to fill completely; thus, a target response Time (RT) of 800 ms was cued during GG trials (Figure 1B). A subset of partial-stop trials was included to assess selective stopping. During partial-stop trials, either the left (stop-left go-right: SG) or right (go-left stop-right: GS) indicator automatically stopped filling before the target. The objective of partial-stop trials was to withhold the response on the stopped side while still responding in time with the target on the non-stopped side (hereby referred to as stop-hand and respond-hand, respectively). The stop-signal delay (SSD) was initially set so that the filling ceased 250 ms prior to the target and was then adjusted in steps of 28 ms (∼4 frames) across partial-stop trial types independently. The SSD was increased (i.e., less time before target) after successful stopping and decreased (i.e., more time before target) after unsuccessful stopping to obtain an average stopping success of ∼50%. Points were awarded based on RT or stop success for each response and were signalled during the intertrial interval (1 – 2 s) by changing the color of the target lines to encourage accurate responding (green: < 25 ms or successful stop; yellow: 26 – 50 ms; orange: 51 – 75 ms; red: > 75 ms or failed stop).

Cues for stopping certainty were presented on every trial for 2 s before trial onset. Stop cues were embedded into the target line color. A black target line indicated a no-stop (0%) chance. Cyan and magenta indicated a low-(33%) or high-(66%) stop certainty (color counterbalanced across participants). Cues were presented over five primary trial types (Figure 1C). A no-stop cue was presented for both hands to assess responses without an expectation of stopping (certain-go trials). A low-stop cue was presented for both hands to assess reactive selective stopping (uninformative trials). A low-stop or high-stop cue was presented for only one hand, while a no-stop cue was presented for the other to assess proactive selective stopping (low-informative and high-informative trials, respectively). GS and SG partial-stop trials were equally distributed for all stop cues. Finally, no target lines were presented in a small number of trials to assess TMS without the influence of response preparation (rest_in-task_ trials).

The experiment was split into behavioral and TMS sessions (mean separation 6.6 days). The behavioral session was always completed first. Initial SSD values for the TMS session were set to the participant’s averages obtained during the behavioral session. Task instructions were given at the start of each session. Participants completed a practice block of certain-go trials and then uninformative trials for familiarisation. Participants were informed that their primary goal was to earn as many points as possible. The block and total score were updated and displayed at the end of each block. The task protocol for each session consisted of 690 trials, split into 15 blocks of 46 trials with a random trial order (see Table 1 for detailed trial numbers). On average, the behavioral and TMS sessions lasted 90 min and 150 min, respectively.

**Table 1:** Trial numbers during anticipatory response inhibition task.

| Cue type | Trial type |  |  |  |
| --- | --- | --- | --- | --- |
|  | GG | SG | GS | Rest |
| Behavioral session |  |  |  |  |
| certain-go | 80 | - | - | - |
| uninformative | 160 | 40 | 40 | - |
| low-informative | 160 | 40 | 40 | - |
| high-informative | 40 | 40 | 40 | - |
| rest<sub>in-task</sub> | - | - | - | 10 |
| TMS session (non-stimulated) |  |  |  |  |
| certain-go | 20 | - | - | - |
| uninformative | 40 | 10 | 10 | - |
| low-informative | 40 | 10 | 10 | - |
| high-informative | 10 | 10 | 10 | - |
| rest<sub>in-task</sub> | - | - | - | 16 |
| TMS session (stimulated) |  |  |  |  |
| certain-go | 72 | - | - | - |
| uninformative | 48 | 12 | 12 | - |
| low-informative | 96 | 24 | 24 | - |
| high-informative | 48 | 48 | 48 | - |
| rest<sub>in-task</sub> | - | - | - | 72 |

Number of go-left go-right (GG), stop-left go-right (SG), go-left stop-right (GS), and rest trials across cue types for behavioral and transcranial magnetic stimulation (TMS) sessions. Nonconditioned, interhemispheric facilitation, and interhemispheric inhibition TMS protocols were equally distributed across stimulated trials.

### Electromyography

Surface EMG was collected from the task-relevant left and right FDI using Ag-AgCl surface electrodes (CONMED) arranged in a belly-tendon montage (Figure 1A). The activity of task-irrelevant left abductor pollicis brevis (APB) was also recorded during the TMS session. A shared ground electrode was positioned on the posterior surface of the left hand. EMG activity was amplified (×1000), bandpass filtered (10 – 1000 Hz), and sampled at 2000 Hz with a CED interface system (MICRO1401mkII; Cambridge Electronic Design). EMG collection was recorded from trial onset (−500 to 1500 ms) during the task and stimulation onset (−150 to 850 ms) during TMS at rest for later offline analysis.

### Transcranial magnetic stimulation

Two MagStim 200^2^ stimulators (The Magstim Company Ltd, Whitland) were used to deliver test and conditioning pulses through 70 mm and 50 mm figure-of-eight coils (100 µs pulse width, monophasic waveform) positioned over right and left M1 respectively. The coils were held by separate experimenters and oriented to induce a posterior-to-anterior flowing current in the underlying cortical tissue. The optimal coil position for eliciting a motor evoked potential (MEP) in the left and then right FDI was assessed and marked on the scalp.

Motor thresholds were determined using a maximum-likelihood parameter estimation by sequential testing strategy (Awiszus, 2011). Rest motor threshold was determined for right FDI and defined as the minimum stimulation intensity required to elicit an MEP amplitude of at least 0.05 mV. Active motor threshold was also determined for right FDI and defined as the minimum stimulation intensity required to elicit an MEP amplitude of at least 0.2 mV during low-intensity voluntary contraction. The TS intensity was determined for left FDI and defined as the minimum stimulation intensity required to elicit an MEP amplitude of at least 0.5 mV at rest. For IHF, the interstimulus interval and CS intensity were set to 6 ms and 60% active motor threshold, respectively (Baumer et al., 2006). For IHI, the interstimulus interval and CS intensity were set to 40 ms and 130% rest motor threshold, respectively (Harris-Love et al., 2007).

TMS was performed prior to and during the task. A total of 24 nonconditioned (i.e., TS only), 24 IHF, and 24 IHI trials were collected for each condition. TMS measures before the task (rest_pre-task_) were collected over 2 blocks of 36 trials. Stimulated trials were intermixed with non-stimulated trials during standard blocks in the TMS session. The TS during stimulated trials was delivered 550 ms after trial onset to avoid contamination by EMG (Cirillo et al., 2018). The Arduino Leonardo response box controlled in-task stimulation timing to ensure synchronisation between task software and TMS equipment. The SSD during stimulated trials was fixed to 550 ms so that stimulation never occurred after a stop signal. Thus, modulation by proactive response inhibition could be isolated from reactive processes.

### Dependent measures

Data were processed using custom scripts in MATLAB (R2021b, v9.11; The MathWorks).

#### Behavioral data

Task data were analyzed from the behavioral session only to avoid the influence of TMS on performance. Responses were coded as errors and removed from further analyses if RTs were more than 400 ms from the target (average 0.27% of trials). Relative RT (RT_rel_) was calculated by subtracting the target time from the mean response time for each hand and trial type. Negative and positive RT_rel_ values indicate early and late responses, respectively. Response delay effect (RDE) provides a measure of proactive response inhibition at a behavioral level (i.e., response slowing). RDE was calculated by subtracting RT_rel_ during certain-go trials from RT_rel_ of the cued hand for each stop-cue type. Stopping-interference provides a measure of stopping selectivity at the behavioral level, where greater values indicate larger interference. The stopping-interference effect was calculated by subtracting the mean RT_rel_ during go trials from the mean RT_rel_ of the respond-hand during successful partial-stop trials. Mean SSD was also calculated for each stop-cue type and made relative to the target, where values closer to 0 indicate less time for stopping.

#### EMG data

EMG data were pre-processed as per Raud et al. (2022). Data epochs were bandpass filtered (20 – 250 Hz) using a second-order Butterworth filter and then resampled to 500 Hz. Filtered data were smoothed by taking root mean square over a 50 ms sliding window and then normalized to baseline (200 – 400 ms after trial onset). Lastly, data epochs were z-scored for each hand and participant separately. An EMG burst was identified if activity above threshold (1.2 z) was found between 0.4 and 1.2 s after trial onset. Burst onset and offset were determined by taking the time of the first data point in a consecutive group of five (i.e., 10 ms) that were below the threshold when working backwards (onset) and forwards (offset) from the peak. The peak rate of rise of the EMG burst (i.e., z/s) was determined as the maximum value of the differentiated signal between burst onset and peak.

EMG dependent measures included burst-onset, burst-amplitude, burst-rise from successful partial-stop trials where an EMG burst was present in both the respond-hand and stop-hand (average 33.9% of partial-stop trials). Cancel time was calculated by subtracting SSD from EMG burst offset in the stop-hand trial-by-trial (Jana et al., 2020). In this case, smaller cancel time values reflect faster stopping or faster cessation of going processes relative to stop signal onset (Raud et al., 2022). Δburst-onset was calculated by subtracting the EMG burst onset of the stop-hand from the respond-hand. Δburst-onset is an EMG proxy of stopping-interference that can be quantified on a trial-by-trial basis, where values greater than 0 indicate nonselective stopping (Raud et al., 2020).

#### TMS data

Peak-to-peak MEP amplitude was calculated for the left FDI and APB between 10 and 50 ms after stimulation. MEPs were excluded when pre-trigger root mean square EMG activity exceeded 20 µV in a -100 to -50 ms pre-trigger window (average 1.15% of trials). The top and bottom 10% of MEP amplitudes were trimmed for each condition before calculating the mean MEP amplitude (Wilcox, 2010), provided that at least 10 MEPs were available. Corticomotor excitability (CME) was determined as the mean of NC MEP amplitude for each condition. The magnitude of IHF and IHI were calculated for each condition as follows:

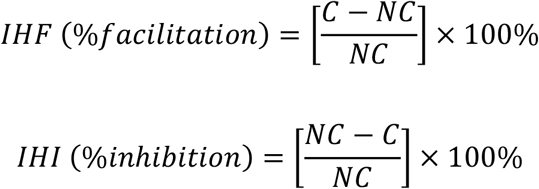

where C and NC refer to each participant’s mean conditioned and nonconditioned MEP amplitude. For ΔCME, ΔIHF, and ΔIHI were computed to determine modulation by proactive cueing by subtracting the mean value during certain-go trials from the mean value observed for each stop-cue type. Values greater than 0 represent the upregulation of a particular measure relative to a certain-go response context.

### Statistical analyses

Data were analyzed with Bayesian repeated-measures analyses of variance (ANOVA) using the BayesFactor package (Morey & Rouder, 2021) in R software (version 4.1.2; R Core Team, 2021). All models included random slopes and were fitted across 100,000 iterations with participant modelled as a random intercept (van den Bergh et al., 2022). Normality of data and model-averaged residual plots were visually inspected before ANOVA (van den Bergh et al., 2020). Logarithmic transformations were used for non-normal data. Interaction effects were determined by comparing models with the interaction term against matched models without the term (van den Bergh et al., 2020). Evidence for main effects and interactions were determined using Bayes factor in favor of the alternative hypothesis (BF_10_ ± percent error), where values greater than 1 indicate support for the alternative hypothesis and values less than 1 support the null hypothesis. The strength of evidence was determined using a standard BF_10_ classification table (BF10 < 0.3: moderate evidence for null hypothesis; 0.3 ≤ BF10 ≤ 3: inconclusive evidence; BF10 > 3: moderate evidence for alternative hypothesis; van Doorn et al., 2021). Post-hoc pairwise comparisons were performed using Bayesian paired t-tests when a main effect or interaction was found. Corrected posterior odds (*O*_post._) were calculated by multiplying the uncorrected BF_10_ by the adjusted prior odds (van den Bergh et al., 2020). Prior odds were adjusted using the Westfall multiple comparisons approach (Westfall, 1997). Data are presented as non-transformed means ± SD unless otherwise specified.

Behavioral and electromyography data were assessed using two-way ANOVAs with the factors of Cue (uninformative, low-informative, high-informative) and Hand (left, right) to determine the influence of stopping certainty on proactive response slowing (RDE), stopping speed (SSD, cancel time), and stopping selectivity (stopping-interference, Δburst-onset). EMG burst onset, amplitude and rise were modelled with one-way ANOVAs with the factor of Cue (uninformative, low-informative, high-informative) to determine how proactive response inhibition affected the respond-hand and stop-hand. In this case, the analyses were performed separately for the respond-hand and stop-hand during successful partial-stop trials where an EMG burst was observed in both hands. CME, IHF, and IHI during rest and response preparation were analyzed with two-way ANOVAs with the factor of Context (rest_pre-task_, rest_in-task_, certain-go) and Muscle (FDI, APB) to determine if TMS measures were modulated by context. ΔCME, ΔIHF, and ΔIHI were modelled with two-way ANOVAs with the factors of Cue (low-informative, high-informative) and Muscle (FDI, APB) to determine modulation by stopping certainty. In this case, the analyses were performed when the left side was cued to stop (i.e., stop-left cues) and cued to go (i.e., stop-right cues).

## Results

### Behavioral results

Behavioral data are shown in Table 2 and Figure 2. For RDE, there was a main effect of Cue (BF_10_ = 7.65 × 10^14^ ± 0.01%). RDE for uninformative trials (9.1 ± 11.3 ms) was less than low-informative (22.8 ± 17.9 ms; *O*_post._ = 6.36 × 10^4^) and high-informative trials (50.4 ± 41.4 ms; *O*_post._ = 2.10 × 10^6^), which also differed from each other (*O*_post._ = 95.04). There was a null main effect of Hand (BF_10_ = 0.18 ± 0.02%) and a Cue × Hand interaction (BF_10_ = 0.14 ± 0.03%). For SSD, there was a main effect of Cue (BF_10_ = 2.42 × 10^33^ ± 0.01%). SSD for uninformative trials (−248.0 ± 25.9 ms) was less than low-informative (−216.1 ± 25.7 ms; *O*_post._ = 1.71 × 10^6^) and high-informative trials (−171.7 ± 30.6 ms; *O*_post._ = 3.21 × 10^18^), which also differed from each other (*O*_post._ = 1.84 × 10^9^). There was a null main effect of Hand (BF_10_ = 0.18 ± 0.01%) while the Cue × Hand interaction was inconclusive (BF_10_ = 0.52 ± 0.02%). Responses slowed and the required time for stopping decreased as stopping certainty increased.

**Table 2.** Results for go and partial-stop trials during the behavioral session.

| Cue | Left RT<sub>rel.</sub>, ms | Right RT<sub>rel.</sub>, ms | Stop success, % |
| --- | --- | --- | --- |
| certain-go | 6.5 (38.7) | 5.8 (37.2) | - |
| uninformative | 15.2 (39.4) | 15.3 (38.6) | 49.9 (2.1) |
| low-informative (left) | 28.9 (42.6) | 5.5 (39.0) | 51.7 (2.6) |
| low-informative (right) | 3.0 (40.3) | 29.0 (39.9) | 53.0 (3.7) |
| high-informative (left) | 56.4 (52.6) | 2.4 (42.8) | 63.0 (12.0) |
| high-informative (right) | -3.8 (40.0) | 56.7 (60.4) | 65.8 (9.1) |

Values reflect mean (SD). Relative response time (RT<sub>rel.</sub>) is calculated by subtracting the target time (800 ms) from mean absolute response time. Negative and positive values indicate early and late responses, respectively. Stop success reflects percentage of trials where the response was successfully withheld on the stop-hand.

**Figure 2:**
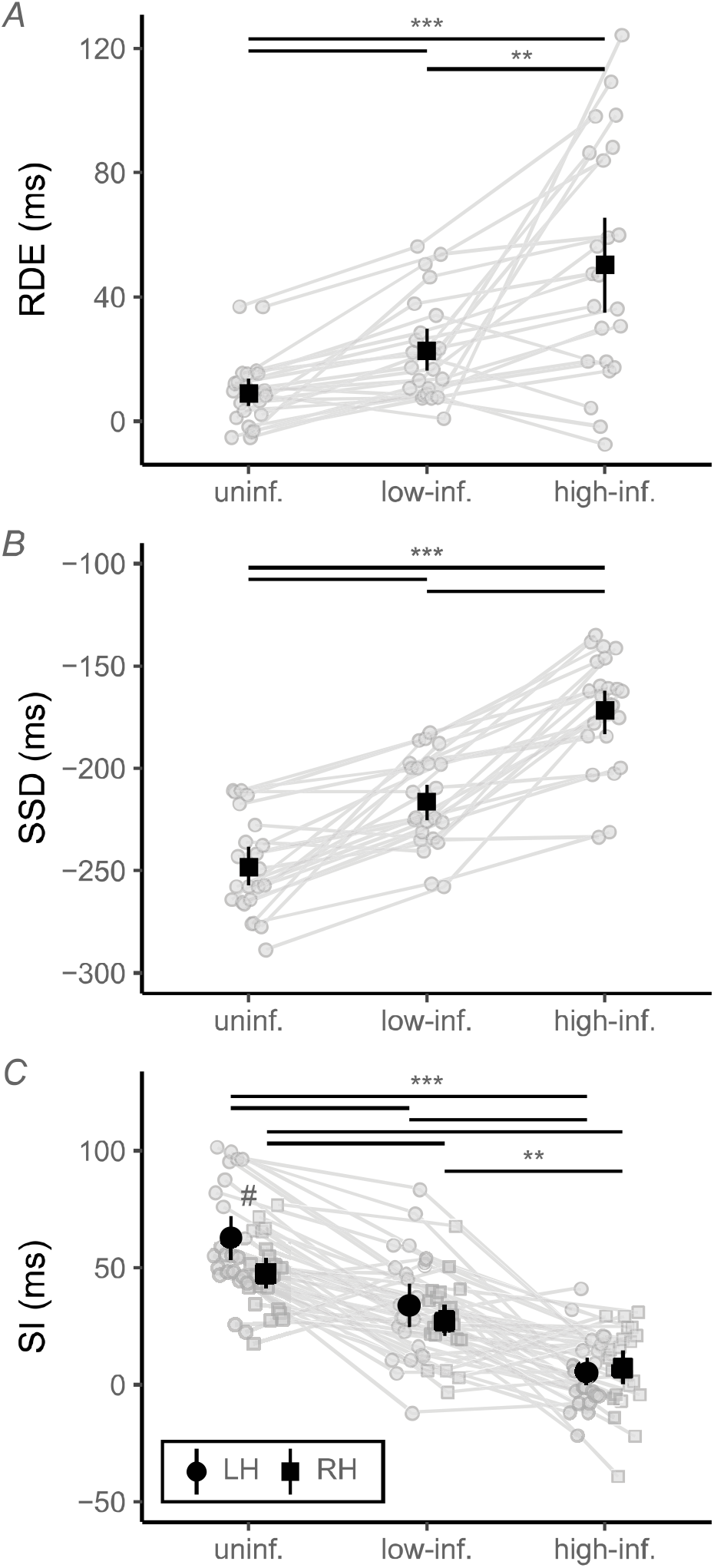
Behavioral measures from partial-stop trials. *A*: Response delay effect (RDE) collapsed across left hand (LH) and right hand (RH). RDE is calculated as the difference in response times between stop-cued and certain go trials, where values greater than 0 reflect response slowing. *B*: Stop-signal delay (SSD) collapsed across LH and RH. SSD is expressed as time relative to target response time, where values closer to 0 reflect stop signals closer to the target. *C*: Stopping-interference (SI) calculated as the difference between mean response times during go trials from the response time of the respond-hand during successful partial-stop trials. Point ranges represent means with 95% bootstrap confidence intervals. Posterior odds: ** > 10, *** > 100. # Posterior odds > 3 for LH versus RH.

For the stopping-interference effect, there was a moderate Cue × Hand interaction (BF_10_ = 4.00 ± 0.04%). Left hand stopping-interference for uninformative trials (63.4 ± 23.9 ms) was less than low-informative (34.5 ± 23.5 ms; *O*_post._ = 2.46 × 10^3^) and high-informative trials (5.7 ± 15.5 ms; *O*_post._ = 6.35 × 10^6^), which also differed from each other (*O*_post._ = 1.73 × 10^3^). Right hand stopping-interference for uninformative trials (47.7 ± 15.5 ms) was less than low-informative (27.4 ± 15.8 ms; *O*_post._ = 442.32) and high-informative trials (7.7 ± 17.9 ms; *O*_post._ = 2.49 × 10^4^), which also differed from each other (*O*_post._ = 17.31). There was an asynchrony in stopping-interference between the left and right hand for uninformative (*O*_post._ = 24.67) but not low-informative (*O*_post._ = 0.86) or high-informative (*O*_post._ = 0.24) trials. One-sample t-tests against 0 indicated a stopping-interference effect for uninformative (*O*_post._ = 9.92 × 10^9^) and low-informative (*O*_post._ = 3.07 × 10^5^) partial-stop trials, but inconclusive evidence for high-informative partial-stop trials (*O*_post._ = 2.35). The selectivity of stopping improved at a behavioral level as stopping certainty increased.

### Electromyography

One participant was excluded from EMG analyses due to a technical issue during data collection, leaving 22 participants available for EMG analyses. EMG data are shown in Figure 3. For cancel time, there was a main effect of Cue (BF_10_ = 4.54 × 10^17^ ± 0.04%).

**Figure 3:**
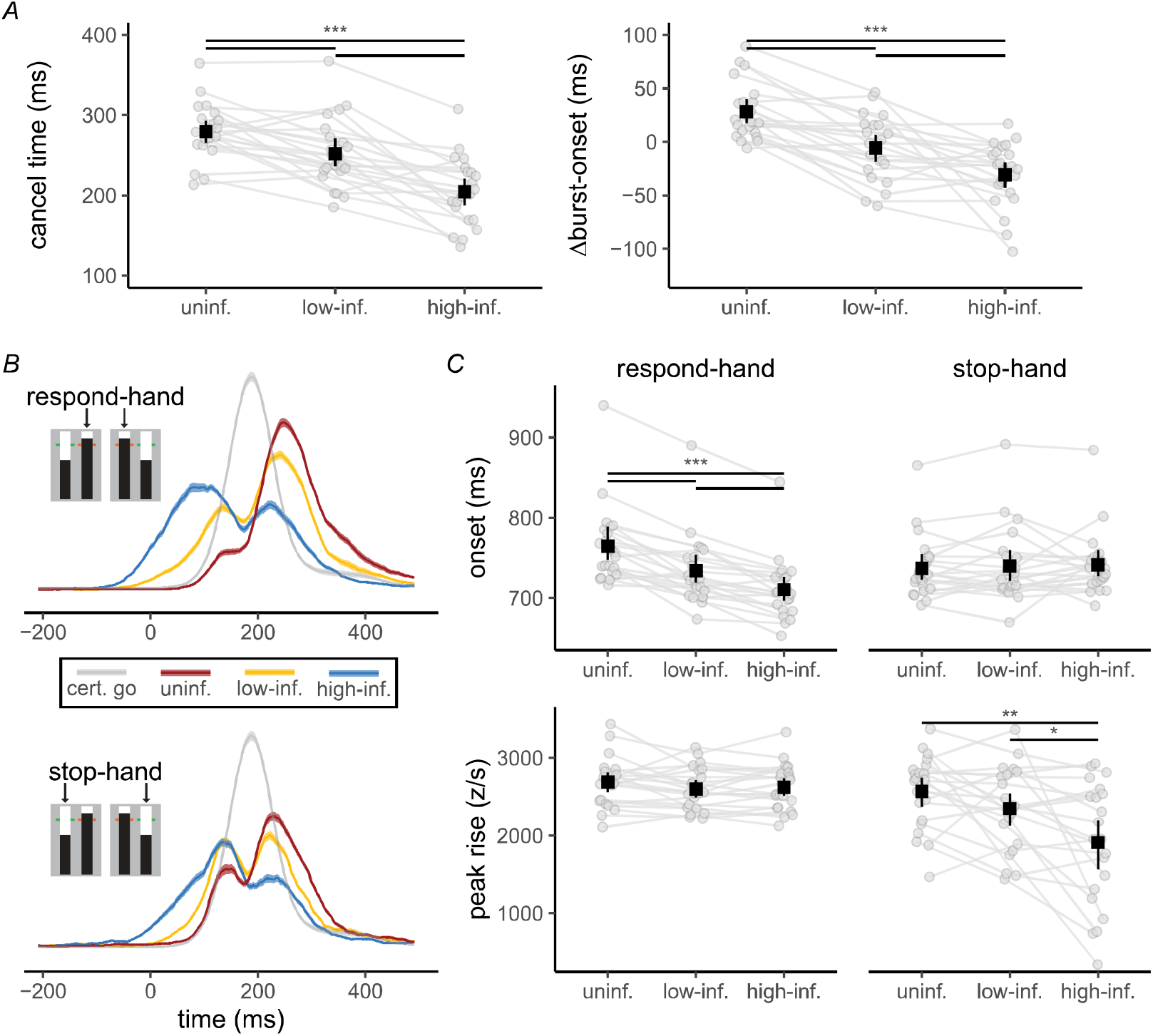
Electromyography (EMG) characteristics during successful partial-stop trials where a burst was observed in both the stop-hand and respond-hand. *A*: Modulation of stopping speed (cancel time) and selectivity (Δburst-onset) by stopping certainty (uninformative, low-informative, high-informative). Cancel time is calculated as the difference in time between stop-signal and EMG burst-offset in the stop-hand, where greater values indicate slower stopping. Δburst-onset is calculated as difference in onset time between the respond-hand and stop-hand, where values above 0 indicate within-trial stopping-interference. *B*: Stop-locked grand-average EMG traces from successful partial-stop trials. Values reflect means with 95% confidence interval bands. Certain go trials are time locked to 550 ms (equivalent to average stop time for uninformative partial-stop trials). *C*: Modulation of EMG burst-onset and burst-rise by stopping certainty. Point ranges represent means with 95% bootstrap confidence intervals. Posterior odds: * > 3, ** > 10, *** > 100.

Cancel time for uninformative trials (279.4 ± 38.4 ms) was greater than low-informative (251.8 ± 48.8 ms; *O*_post._ = 174.95) and high-informative trials (204.3 ± 46.9 ms; *O*_post._ = 9.05 × 10^11^), which also differed from each other (*O*_post._ = 3.21 × 10^4^). There was a null main effect of Hand (BF_10_ = 0.19 ± 0.01%) and a Cue × Hand interaction (BF_10_ = 0.14 ± 0.04%). For Δburst-onset, there was a main effect of Cue (BF_10_ = 8.33 × 10^13^ ± 0.03%). Δburst-onset for uninformative trials (28.1 ± 30.1 ms) was greater than low-informative (−5.7 ± 35.1 ms; *O*_post._ = 3.80 × 10^5^) and high-informative trials (−30.7 ± 37.6 ms; *O*_post._ = 4.05 × 10^6^), which also differed from each other (*O*_post._ = 23.39). There was a null main effect of Hand (BF_10_ = 0.20 ± 0.03%) and a Cue × Hand interaction (BF_10_ = 0.28 ± 0.04%). There was no observable interference in EMG when informative stop-cues were presented.

Grand average EMG bursts from the respond-hand and stop-hand during successful partial-stop trials are shown in Figure 3B. For the respond-hand, there was a main effect of Cue on burst-onset (BF_10_ = 2.06 × 10^9^ ± 0.01%). Burst-onset occurred earlier for high (710.4 ± 38.6 ms) compared to low-informative (734.2 ± 43.7 ms; *O*_post._ = 1.92 × 10^3^) and uninformative partial-stop trials (764.8 ± 48.9 ms; *O*_post._ = 5.27 × 10^4^), which also differed from each other (*O*_post._ = 2.16 × 10^3^). There was a main effect of Cue on burst-amplitude (BF_10_ = 5.02 ± 0.03%), however, post-hoc tests did not provide conclusive evidence (all *O*_post._ < 3). The main effect of Cue on burst-rise was inconclusive (BF_10_ = 0.95 ± 0.03%). For the stop-hand, there was a null main effect of Cue on burst-onset (BF_10_ = 0.18 ± 0.03%) and inconclusive evidence for burst-amplitude (BF_10_ = 0.33 ± 0.04%). There was a main effect of Cue on burst-rise (BF_10_ = 627.07 ± 0.01%). Burst-rise was smaller for high-informative trials (1909.7.9 ± 772.6 z/s) compared to low-informative (2346.3 ± 533.1 z/s; *O*_post._ = 3.55) and uninformative partial-stop trials (2568.9 ± 464.7 z/s; *O*_post._ = 61.26), which did not differ from each other (*O*_post._ = 0.63). Successful partial-stop trials were marked by a smaller EMG burst-onset and burst-rise in the respond-hand and stop-hand respectively (Figure 3C).

### TMS results

Five participants were excluded from the TMS session due to high resting FDI resting motor thresholds which caused coil overheating during task blocks. The high motor thresholds were likely due to the small coil size required for dual-hemisphere stimulation. After exclusions, data from 17 participants were available for TMS analyses. The main findings did not change if participants with incomplete datasets were excluded from behavioral analyses. The TS intensity for the remaining participants was 53.5 ± 10.6% of maximum stimulator output (MSO). The CS intensity for the IHF and IHI protocols were 29.6 ± 6.2% and 75.7 ± 16.9% MSO, respectively. On average, 23.7 ± 0.9 trials (range 16 – 24 trials) were available for each TMS trial type. Figure 4A shows representative MEPs in EMG traces from one participant.

**Figure 4:**
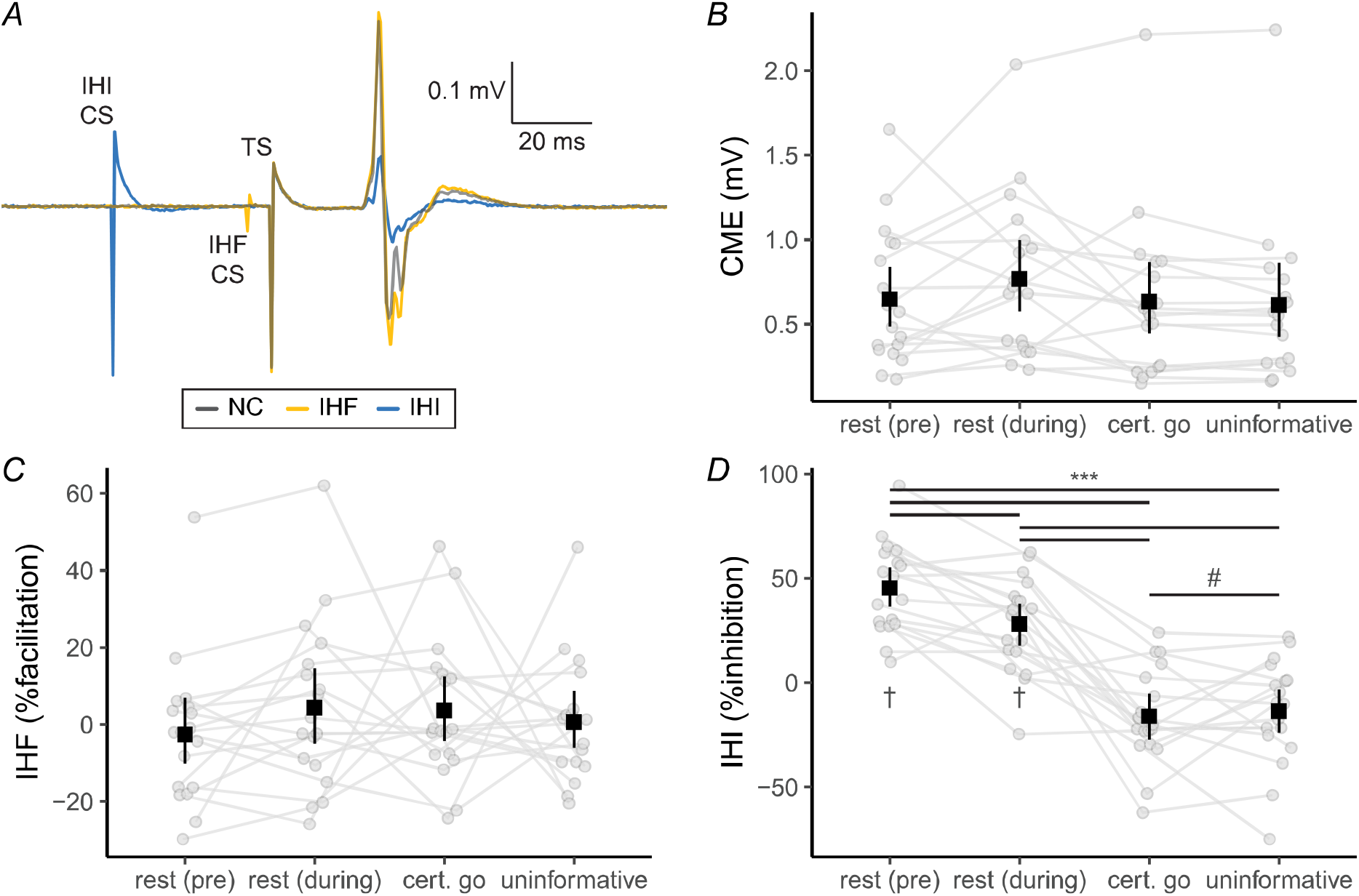
Transcranial magnetic stimulation was used to examine corticomotor excitability (CME), interhemispheric facilitation (IHF) and interhemispheric inhibition (IHI). *A*: Electromyography traces with motor evoked potentials (MEPs) in the task-relevant left first dorsal interosseous muscle during pre-task rest. Each trace is the average of 24 trials from a representative participant. For IHF, a subthreshold conditioning stimulus (CS) was delivered 6 ms prior to the test stimulus (TS). For IHI, a suprathreshold CS was delivered 40 ms prior to the TS. *B*: Corticomotor excitability (CME) calculated as mean peak-to-peak motor MEP amplitude. *C*: IHF calculated as percent facilitation, where greater values indicate a larger conditioned relative to nonconditioned MEP amplitude (i.e., more facilitation). *C*: IHI calculated as percent inhibition, where greater values indicate a smaller conditioned relative to nonconditioned MEP amplitude (i.e., more inhibition). † Posterior odds > 3 from one-sample t-test against 0. Point ranges represent means with 95% bootstrap confidence intervals. *** Posterior odds > 100. # Posterior odds < 0.03

#### Context-dependent modulation of TMS-derived measures

TMS data are shown in Figure 4. For CME, there was a main effect of Muscle (BF_10_ = 5.55 × 10^6^ ± 0.01%). CME was less in FDI (0.66 ± 0.41 mV) compared to APB (0.67 ± 0.89 mV; *O*_post._ = 10.17) across response contexts. There was a null Context x Muscle interaction (BF = 0.16 ± 0.00%) while the main effect of Context was inconclusive (BF_10_ = 1.69 ± 0.00%). For IHF, there was a null main effect of Muscle (BF_10_ = 0.20 ± 0.01%) and Context (BF_10_ = 0.21 ± 0.01%), as well as a null Context × Muscle interaction (BF_10_ = 0.09 ± 0.02%). One-sample t-tests against 0 indicated no IHF across response contexts (all *O*_post._ < 0.3).

For IHI, there was a null main effect of Muscle (BF_10_ = 0.19 ± 0.01%) and a null Context × Muscle interaction (BF_10_ = 0.19 ± 0.01%). There was a main effect of Context (BF_10_ = 3.59 × 10^29^ ± 0.02%). IHI during rest_pre-task_ (45.4 ± 22.9%) was greater than rest_in-task_ (28.1 ± 23.3%; *O*_post._ = 111.23), certain-go (−16.1 ± 27.5%; *O*_post._ = 5.01 × 10^9^), and uninformative contexts (−13.7 ± 28.0%; *O*_post._ = 3.97 × 10^10^). Rest_in-task_ was also greater than certain-go (*O*_post._ = 6.46 × 10^5^) and uninformative contexts (*O*_post._ = 1.18 × 10^6^). There was no difference between certain-go and uninformative contexts (*O*_post._ = 0.05). One-sample t-tests against 0 provided indicated IHI during rest_pre-task_ (*O*_post._ = 6.35 × 10^4^) and rest_in-task_ (*O*_post._ = 170.62) contexts, while IHI during certain-go (*O*_post._ = 2.55) and uninformative contexts (*O*_post._ = 0.81) was inconclusive. A post-hoc one-way ANOVA on average MEP amplitudes elicited in the right FDI from the CS during IHI trials indicated a null main effect of Cue (BF_10_ = 0.13 ± 0.02%). IHI was modulated by context for task-relevant and irrelevant muscles.

#### Cue-dependent modulation of CME, IHF and IHI toward the left hand

Cue-dependent modulation of motor evoked potentials are shown in Figure 5. For ΔCME there was a main effect of Muscle (BF_10_ = 15.89 ± 0.01%) when the left side was cued to stop. ΔCME was larger in FDI (−0.07 ± 0.14 mV) compared to APB (0.00 ± 0.09 mV; *O*_post._ = 5.30). There was a null main effect of Cue (BF_10_ = 0.28 ± 0.00%) while the Cue × Hand interaction was inconclusive (BF_10_ = 0.49 ± 0.02%). When the left side was cued to Go (stop-right) the main effect of Muscle (BF_10_ = 0.78 ± 0.01%) and Cue (BF_10_ = 0.41 ± 0.01%) were inconclusive. There was a Cue × Hand interaction (BF_10_ = 5.23 ± 0.02%), however, post-hoc comparisons were inconclusive (all *O*_post._ < 3).

**Figure 5:**
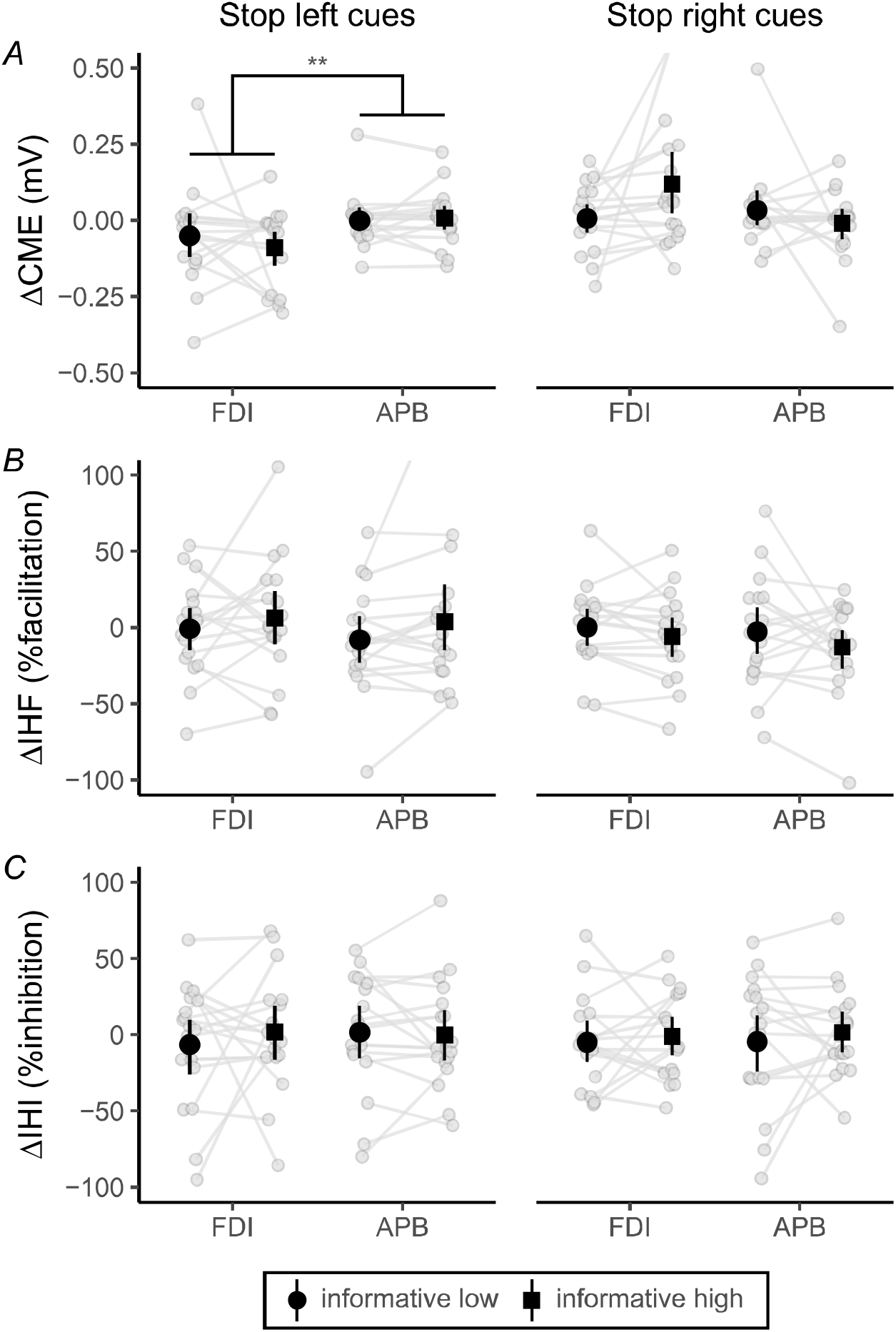
Modulation of transcranial magnetic stimulation measures by informative partial-stop cues in the task-relevant left first dorsal interosseous (FDI) and task-irrelevant abductor pollicis brevis (APB) muscles. All measures have been calculated as the difference from certain go trials, where positive and negative values indicate upregulation and downregulation, respectively. *A*: Corticomotor excitability (CME) calculated as mean peak-to-peak motor evoked potential (MEP) amplitude. *B*: Interhemispheric facilitation (IHF) calculated as percent facilitation, where greater values indicate larger conditioned relative to nonconditioned MEP amplitude (i.e., more facilitation). *C*: Interhemispheric inhibition (IHI) calculated as percent inhibition, where greater values indicate smaller conditioned relative to nonconditioned MEP amplitude (i.e., more inhibition). Point ranges represent means with 95% bootstrap confidence intervals. ** Posterior odds > 10

For ΔIHF, when cued to stop the main effect of Muscle (BF_10_ = 0.32 ± 0.01%), Cue (BF_10_ = 0.78 ± 0.01%) and a Cue × Hand interaction (BF_10_ = 0.33 ± 0.02%) were inconclusive. When cued to Go (stop-right), the main effect of Muscle (BF_10_ = 0.41 ± 0.01%), Cue (BF_10_ = 1.04 ± 0.01%) and a Cue × Hand interaction (BF_10_ = 0.34 ± 0.02%) were inconclusive. For ΔIHI when cued to stop, there was a null main effect of Muscle (BF_10_ = 0.28 ± 0.04%) and Cue (BF_10_ = 0.28 ± 0.02%), but the Cue × Hand interaction was inconclusive (BF_10_ = 0.43 ± 0.03%). For ΔIHI when cued to go (stop-right cues), there was a null main effect of Muscle (BF_10_ = 0.27 ± 0.06%). The main effect of Cue (BF_10_ = 0.52 ± 0.16%) and Cue × Hand interaction (BF_10_ = 0.35 ± 0.04%) were inconclusive.

## Discussion

The present study provides new insights into M1-M1 interhemispheric influences during selective stopping with proactive cueing. In support of the first hypothesis, response times were prolonged and stopping-interference was less as stopping certainty increased. The second hypothesis was partly supported by a release of IHI from rest to in-task contexts in both the task-relevant and task-irrelevant effector. The third hypothesis was not supported since there was no evidence for effector-specific modulation of IHF by selective stopping cues. Overall, the findings indicate that interhemispheric M1-M1 channels are involved in setting inhibitory tone across response contexts but are not directly involved in proactive response inhibition during selective stopping.

### Greater selectivity of stopping with proactive cueing

The difference in response times between certain-go and stop-cued contexts (RDE) increased progressively with stopping certainty. Response slowing was specific to the cued hand and likely reflected the behavioral manifestation of proactive response inhibition to improve stopping (Verbruggen & Logan, 2009). Since response delays tended to be less than 50 ms, this indicates that responses were slowed rather than omitted entirely (Verbruggen et al., 2013). Stopping performance improved from uninformative to informative partial-stop trials as evident by shorter SSDs. Cancel times during successful partial-stop trials were also shorter with greater stopping certainty. Cancel times were calculated on a trial-by-trial basis and reflect the time taken to inhibit or disengage excitatory processes in the stop-hand (Raud et al., 2022). The shorter SSDs and cancel times indicate that stopping was faster during informative partial-stop trials. Interestingly, the cancel time in the reactive (uninformative) context was ∼100 ms longer than reported during unimanual stopping (Jana et al., 2020; Raud et al., 2022). Although nonselective stopping was not explicitly assessed, the cancel times in the present study provide evidence of slower stopping in selective compared to nonselective stopping contexts (Smittenaar et al., 2013; c.f., Raud & Huster, 2017).

Stopping selectivity was better with proactive cueing. The stopping-interference effect was smaller during low-informative compared to uninformative partial-stop trials (Cai et al., 2011; Cirillo et al., 2018; Drummond et al., 2018; Lavallee et al., 2014; Majid et al., 2012; Raud & Huster, 2017; Smittenaar et al., 2013). Stopping-interference was smallest during high-informative partial-stop trials and larger in the left hand than the right hand during uninformative but not informative partial-stop trials. A between-hand discrepancy in stopping-interference may result from hand dominance and coupling. The functional coupling account of the stopping-interference effect posits that part of the response delay arises from decoupling the respond-hand from the stop-hand (Wadsley et al., 2022b). Thus, stopping-interference may be larger in the nondominant hand as it is more stringently coupled to the dominant hand than vice-versa (Byblow et al., 2000). The discrepancy may be absent during informative partial-stop trials since proactive response inhibition processes can be directed to the cued hand irrespective of dominance.

No stopping-interference was observed during informative partial-stop trials with a trial-by-trial measure of response delays (Δburst-onset). Indeed, EMG burst-onsets were only delayed in the respond-hand relative to the stop-hand during uninformative partial-stop trials. Greater stopping selectivity during informative partial-stop trials may have occurred through separate adjustments to the planned action in the respond-hand and stop-hand. For the respond-hand, EMG burst-onset but not peak rate of rise decreased with stopping certainty and indicates that an equivalent go process was planned but released earlier (Coxon et al., 2007; Macdonald et al., 2012). In contrast, EMG burst-onset was unchanged, but the peak rate of rise was smaller in the stop-hand with greater stopping certainty. A smaller rate of increase but not different onset is indicative of proactive response inhibition that specifically suppressed the stop-cued hand (MacDonald et al., 2017). Therefore, better selective stopping with proactive cueing may have occurred through suppression of the stop-hand and an earlier release of the go process in the respond-hand. The above findings indicate that stopping can proceed selectively for both hands with proactive cueing.

### Modulation of IHI but not IHF from rest to in-task contexts

There were no observations of IHF across response contexts. We hypothesized that IHF would be elicited within a task-relevant effector during action preparation. However, there was moderate evidence for a null effect of the IHF protocol in both rest and in-task contexts. Reports of M1-M1 facilitation are limited (Baumer et al., 2006) and may be observed at longer interstimulus intervals (Fiori et al., 2017). Alternatively, IHF may be driven by premotor-M1 interactions (Neige et al., 2021). IHF at similarly short interstimulus intervals has been observed during task switching when conditioning the premotor cortex (Mars et al., 2009). In summary, there was no evidence for task-dependent modulation of M1-M1 IHF.

There was a decrease of IHI between resting state and in-task response preparation. A similar decrease in IHI was observed previously with short-interval IHI during a bimanual ARI paradigm (MacDonald et al., 2021) and may reflect disinhibition for action selection. It could be advantageous for IHI to be maintained in task-irrelevant effectors as a potential centre-surround inhibitory mechanism (Hinder et al., 2018). However, in the current study, a release of IHI was also observed in the task-irrelevant APB muscle. This nonspecific release of IHI may reflect a wide aperture of sensorimotor disinhibition and the functional similarity of FDI and APB for many dexterous tasks (Labruna et al., 2019). Interestingly, IHI was present but smaller during in-task rest compared to pre-task rest. It is unlikely that the observed modulation of IHI was the consequence of general shifts in excitability since CME did not differ across rest and in-task contexts. Therefore, IHI is released during response preparation and reduced in contexts that necessitate frequent responding.

IHI was not modulated between cues signaling certain-go and uninformative trials. An absence of upregulation from a pure go context to one where stopping may be required indicates IHI may not be involved in proactive response inhibition. The pattern of IHI modulation contrasts with that observed during within-hemisphere probes of GABA-B receptor-mediated inhibition, which were elevated for uninformative compared to certain-go trials, and positively associated with the magnitude of stopping-interference (Cowie et al., 2016). The discrepancy may be a consequence of the certain-go context being assessed with a trial-by-trial rather than a block-by-block design since GABA-B is insensitive to trial-wise modulation (Cirillo et al., 2018). The pattern of IHI modulation indicates a role in setting general inhibitory tone that is not engaged proactively to aid response inhibition.

Proactive response inhibition was marked by effector-specific CME suppression. Suppression occurred in the task-relevant effector when cued about stopping (informative stop-left cues; Figure 4A). This suppression corroborates previous investigations of proactive selective stopping and indicates that CME suppression may be a general feature of response preparation in contexts that may require stopping (Cai et al., 2011; Raud et al., 2020). An increase in CME was not observed when the left side was cued to go (i.e., stop-right), despite evidence of an interaction driven by high-informative stop cues. Trial-by-trial changes in CME were not supported by concomitant IHI modulation and instead may have been driven through a separate neural mechanism such as the proactive engagement of the indirect cortico-basal-ganglia pathway (Majid et al., 2013). For example, BOLD signal in striatum scaled with stop-signal probability and behavioral slowing in a unimanual ARI task (Vink et al., 2015; Zandbelt & Vink, 2010). In summary, the present findings indicate that IHI was reduced between resting and response preparation states but did not contribute to cue-dependent CME suppression.

## Limitations

The present study has some limitations. TMS was applied at only one time point that corresponded to the typical SSD required to elicit a 50% success rate for partial-stop trials in healthy young adults (Macdonald et al., 2012; Wadsley et al., 2022a). Proactive modulation of IHI or IHF may occur closer or further from the time of responding, or alternatively, it may drive reactive response inhibition (MacDonald et al., 2021). However, a reliable number of stimulated trials could be collected for each measure of interest by limiting the time points for in-task TMS (Goldsworthy et al., 2016). Another limitation relates to the parameterisation of TMS for investigating IHF, particularly in terms of localising the conditioning pulse. The current study examined only M1-M1 interactions. Future studies might investigate cortico-cortical interhemispheric interactions between premotor and M1 regions during selective stopping.

## Conclusion

The present study provides novel insight into M1-M1 influences during proactive response inhibition and selective stopping. Behaviorally, responses were slowed with proactive cueing and led to both faster and more selective stopping (i.e., reduced stopping-interference). Analyses of EMG bursts during successful partial-stop trials indicate that improved stopping selectivity may have occurred through targeted suppression of the stop-hand and an earlier release of the go process in the respond-hand. There was no evidence of IHF across response contexts, whereas IHI was reduced from pre-task rest to in-task contexts. While an effector-specific suppression of CME marked proactive response inhibition, there was no concomitant increase in IHI. These findings indicate that stopping can become more selective with proactive cueing, although cue-related improvements in task performance are unlikely to reflect proactive engagement of facilitatory or inhibitory M1-M1 influences.

## Acknowledgements

The authors thank April Ren and Kelly Tay for their assistance with data collection.

